# Minimizing cell number fluctuations in self-renewing tissues with a stem cell niche

**DOI:** 10.1101/2022.03.10.483777

**Authors:** Rutger N.U. Kok, Sander J. Tans, Jeroen S. van Zon

## Abstract

Self-renewing tissues require that a constant number of proliferating cells is maintained over time. This maintenance can be ensured at the single-cell level or the population level. Maintenance at the population level leads to fluctuations in the number of proliferating cells over time. Often, it is assumed that those fluctuations can be reduced by increasing the number of asymmetric divisions, i.e. divisions where only one of the daughter cells remains proliferative. Here, we study a model of cell proliferation that incorporates a stem cell niche of fixed size, and explicitly model the cells inside and outside the niche. We find that in this model fluctuations are minimized when the difference in growth rate between the niche and the rest of the tissue is maximized and all divisions are symmetric divisions, producing either two proliferating or two non-proliferating daughters. We show that this optimal state leaves visible signatures in clone size distributions and could thus be detected experimentally.

## I. INTRODUCTION

Many adult tissues, such as the mammalian intestinal epithelium and the skin epidermis, undergo constant selfrenewal supported by stem cells [1, 2]. To ensure homeostasis, the number of proliferating cells needs to be kept constant over the lifespan of the organism. Large fluctuations in the number of proliferating cells could lead to disease or even the death of the organism. For that reason, tight regulation of cell proliferation is required [3–5], where every cell division must result in one proliferating and one non-proliferation cell, at least on average.

The balance of proliferation and differentiation can either be maintained strictly at the single-cell level or at the population level [6]. In the first strategy, every cell division produces an asymmetric outcome: one daughter cell remains proliferative and the other daughter ceases proliferation and terminally differentiates. In the population level strategy, the balance between proliferation and terminal differentiation is only maintained on average at the population level [7]. Individual divisions can result in zero, one or two proliferating daughters. This maintenance strategy is called the population-asymmetry model.

Unlike the single-cell level strategy, the population level strategy is inherently stochastic, and therefore potentially prone to fluctuations in the number of proliferating cells. Despite these fluctuations, the population level strategy of self-renewal is found in many stem cell systems [3]. Examples include the mammalian germline, the intestinal and the epidermis[6, 8–11]. In some organs large variations of the number of stem cells can occur, such as for spermatogenic stem cells in murine testes [7]. As such fluctuations are potentially dangerous, in other organs the number of stem cells appears to be tightly controlled, such as in the small intestine [12]. It is unknown how different strategies of stem cell maintenance affect fluctuations in stem cell numbers.

In many stem cell systems, cell proliferation is also organized in space, with stem cell niches that provide a local environment that maintains stem cells in an undifferentiated and proliferating state [7], while cells outside of such stem cell niches eventually cease proliferation and differentiation. As a consequence, the divisions patterns of proliferating cells also likely vary in space, with more divisions generating proliferating cells within the stem cell niche, and more non-proliferating cells without. How such a spatial segregation of proliferation dynamics might impact fluctuations in cell proliferation remains an open question.

Here, we use a theoretical approach to study the impact of different stem cell maintenance strategies, including whether a stem cell niche is present or not, on fluctuations in the number of proliferating cells. In particular, we compare proliferation dynamics in an uniform, unbounded system, i.e. lacking a stem cell niche, and a system with two compartments, a stem cell niche where cells are geared towards proliferation, and a differentiation compartment where cells are biased towards ceasing proliferation. We analytically derive under which conditions the two compartment is stable as well as the corresponding steady-state number of proliferating cells. We then systematically examine how different parameters, such as the fraction of symmetric and asymmetric division, or the size of the stem cell niche, impact the magnitude of fluctuations. We find that in the uniform model fluctuations are minimized when all divisions are asymmetric, strictly generating one proliferating and one non-proliferating daughter. Surprisingly, we find that in the two compartment model, that incorporates a stem cell niche, fluctuations are instead minimized by a very different strategy: when all divisions are strictly symmetric, generating either two proliferating or two nonproliferating daughters. Finally, our simulations show that these different strategies generate distinct clone size distributions, and could thus potentially be differentiated experimentally [5].

## II. RELATED WORK

Theoretical models have advanced our understanding of stem cell behavior. The models have focused on two main questions. First, what is the impact of various niche setups on cell number fluctuations? And second, what is the role of (a)symmetry in cell divisions [4, 5]?

We will first discuss the impact of niche setup on cell number fluctuations. The models used to study fluctuations can roughly be divided in three classes. A first class of models uses a uniform space with two cell types; proliferating and non-proliferating. Klein et al.[15] used such a model to determine the impact of various division strategies on the fluctuations of cell numbers, but only for individual lineages. Sun and Komarova [16] used the same type of model to test the impact of various feedback mechanisms on cell number fluctuations. Those feedback mechanisms required that cells are aware of the current amount of stem cells in the system. However, it is open question whether such feedback mechanisms are present in stem cell systems.

A second class of models uses a single compartment with a fixed number of stem cells, and no other cell types. These models were used to study competition between lineages. Snippert et al.[10] and Lopez-Garcia et al.[9] demonstrated that the intestinal crypt uses neutral competition, where the progeny of one stem cell eventually takes over the entire niche. Ritsma et al.[11] and Corominas-Murtra [17] studied the dependence of lineage survival on cell position within the niche.

Finally, in a third class of models two cell types (stem and non-stem) are distributed over two compartments. In this work, we will use a model of this class. So far, these models have only been used to study the risk of developing cancer, namely by Cannataro et al.[18, 19] and Shahriyari and Komarova [20]. Therefore, the question of how a compartmentalized system affect fluctuations in the number of cells remains open.

The second question is about the impact of division symmetry. The studies that focused on this question so far used a model with uniform space and both proliferating and non-proliferating cells. Klein et al.[15] used this model to infer division symmetry from experimental data of the skin epidermis, and concluded that asymmetric divisions were dominant. Sei et al.[21] used a similar setup for the intestinal crypt, and likewise concluded that asymmetric divisions were dominant. Yang and Komarova [22] used the model for a different purpose: they investigated the impact of division symmetry on fluctuations in the number of proliferating cells. The authors found that their results were mixed. Under some control mechanisms symmetric divisions provide lower fluctuations, but under most control mechanisms asymmetric divisions provided lower fluctuations. Therefore, it remains unclear what impact division symmetry has on the fluctuations in cell numbers.

## III. ONE-COMPARTMENT MODEL

We start our analysis by looking at a model without space. In the next section, this model will be extended to include a niche compartment. We use two cell types: proliferating and non-proliferating. Whether a cell is proliferative or not is decided at the birth of the cell; this fate cannot be changed later. Every proliferating cell will divide *T* hours after the birth of the cell, while non-proliferating cells will never divide. Values of *T* are drawn from a skew-normal distribution [23] with a skewness parameter 6.1, a location of 12.2 and a scale of 5.3. This distribution approximates cell cycle times we recently measured in intestinal organoid crypts [24].

After a division, two daughter cells are created, which can be (**I**) both proliferating, (**II**) both non-proliferating or (**III**) one can be proliferating and the other nonproliferating (Fig. 1a). The chances for these division types to occur are *p, q* and 1 – *p* – *q*, respectively. Division types **I** and **II** are symmetric, while type **III** is asymmetric. The values of *p* and *q* are determined by two parameters. The parameter *ϕ* = *p* + *q* is the chance of a division being symmetric (Fig. 1b), while the other parameter, the growth rate *α* = *p* – *q*, is the average increase in the number of proliferating cells per division (Fig. 1c). One can verify that for *ϕ* = 0 all divisions are required to be of type **III**, while for *α* = 1 all divisions are required to be of type **I**.

**FIG. 1.**
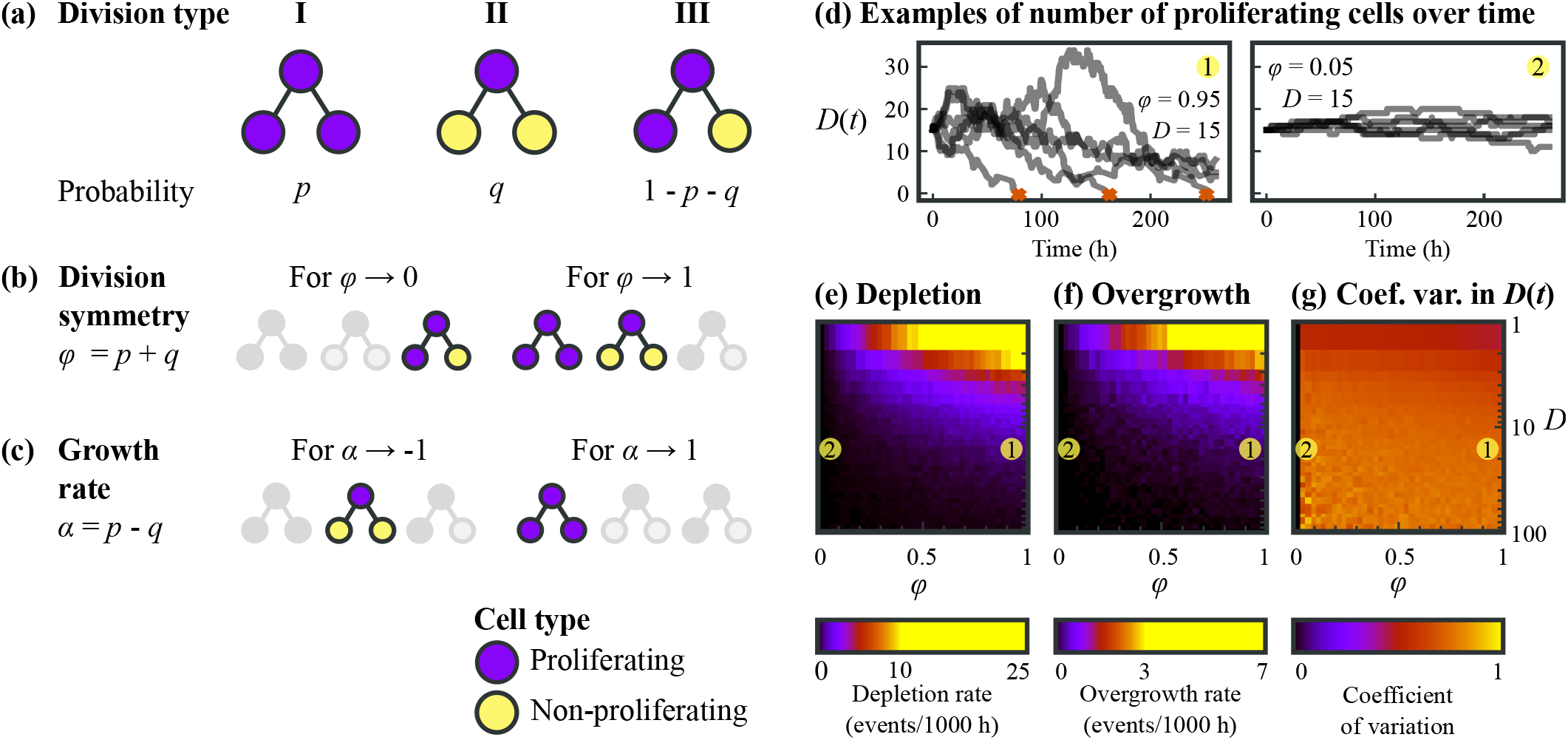
The uniform model. **(a)** All possible division types and their probability. **(b)** Dominant division type for two different division symmetries. **(c)** Dominant division type for two different growth rates. **(d)** Two panels, each showing for six simulations the number of proliferating cells over time. The simulations use *D* = 15 and the given symmetry fraction *ϕ*. **(e)** Depletion rate, the rate of how often the number of proliferating cells drops to zero. **(f)** Overgrowth rate, the rate of how often the number of proliferating cells grew larger than 5 times the initial number of proliferating cells. **(g)** Coefficient of variation of the number of dividing cells.

For homeostasis, in this system it is required that the growth rate *α* = 0. For *α* > 0 the number of proliferating cells would grow exponentially, and for *α* < 0 this number would decrease exponentially. The symmetry fraction *ϕ* can be varied freely, as well as the initial number of proliferating cells *D*. We therefore perform a simulation for different combinations of *D* and *ϕ*, while keeping *α* at 0. The results are displayed in Fig. 1d-g.

In Fig. 1d six simulations of the number of proliferating cells are shown over time for two values of the symmetry fraction *ϕ*. We can observe that a higher fraction of asymmetric divisions (low *ϕ*) provides a system with less fluctuations in the number of proliferating cells. In addition, we see that the system with a high fraction of symmetric divisions (high *ϕ*) is frequently depleted of proliferating cells, while the smaller system with a high fraction of asymmetric divisions (low *ϕ*) remains stable for at least 10 days. In Fig. 1e-g the depletion rate, overgrowth rate and coefficient of variation in the number of proliferating cells are plotted as a function of both the initial number of proliferating cells *D* and the symmetry fraction *ϕ*. Here, depletion is defined as the number of proliferating cells becoming zero, while overgrowth is defined as the number of proliferating cells reaching five times the initial amount. We can see that the depletion and overgrowth rates increase for smaller *D* and *ϕ*, indicating a less stable system. The coefficient of variation remains high irrespective of *D* and *ϕ*, with one exception: the theoretical case where precisely all cell divisions are asymmetric the coefficient is zero.

As a result, in this model the best approach to minimize fluctuations and avoid depletion or overgrowth of proliferating cells would be to have a large number of proliferating cells and to use strictly asymmetric divisions. A low amount of symmetric divisions already results in relatively large fluctuations in the number of proliferating cells.

## IV. TWO-COMPARTMENT MODEL

We wanted to compare the performance of the uniform model, in terms of the impact of fluctuations, to a model that incorporated a stem cell niche. We therefore constructed a different model, that included two compartments, with cell proliferation differing between compartments. One compartment, which we call the niche compartment, can only contain a fixed number of cells. In contrast, the other other compartment, which we call the differentiation compartment, is unbounded in size. This two-compartment model is sketched in Fig. 2.

**FIG. 2.**
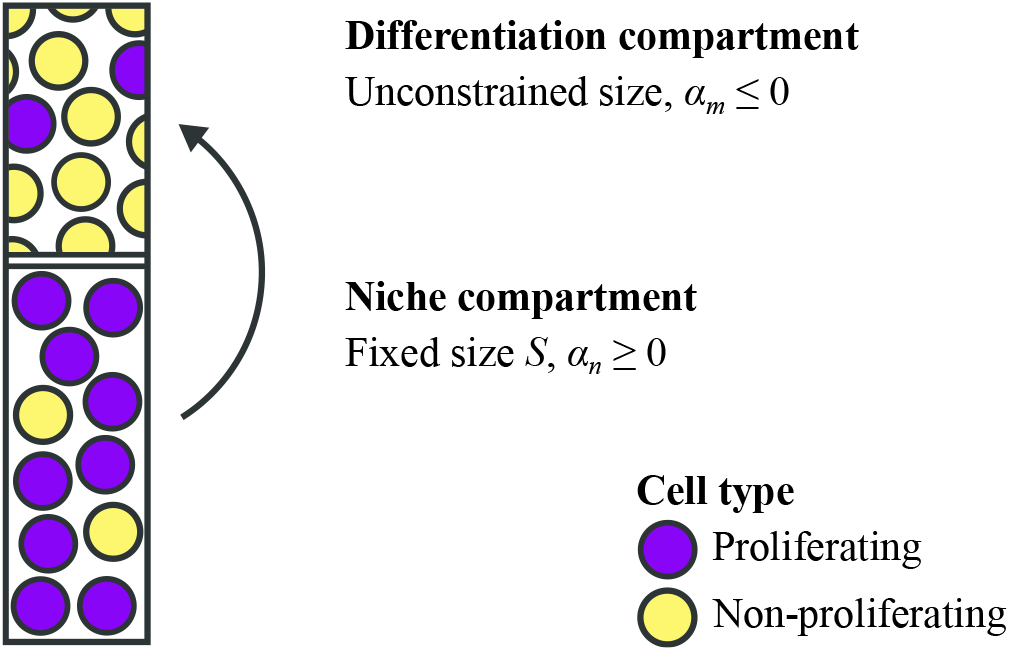
Illustration of the two-compartment model. Two compartments are defined, the niche compartment and the differentiation compartment. In the niche compartment the growth rate *α_n_* ≥ 0 while in the differentiation compartment the growth rate *α_m_* ≤ 0. The fraction of symmetric divisions *ϕ* is equal in both compartments. Upon a division in the niche compartment, one random cell moves from the niche compartment to the differentation compartment.

The niche compartment is set to contain a fixed number of cells in total, denoted as *S*. Of this number, a variable number of cells *N*(*t*) is proliferative, which makes the number of non-proliferating cells equal to *S* – *N*(*t*). For the differentiation compartment we denote the number of proliferating cells as *M* (*t*). As the differentiation compartment is unbounded in size, the number of nonproliferating cells remains unspecified. The total number of proliferating cells over both compartments is defined as *D*(*t*) = *N*(*t*) + *M*(*t*). To keep the niche compartment fixed in size, upon every division in the niche compartment we move one random cell out of the niche compartment into the differentiation compartment.

Cell divisions still follow the rules established in the previous section (Fig. 1a-c). The key difference is that instead of using a single growth rate *α*, both compartments can now have a different growth rate. We define the niche compartment as having a growth rate of *α_n_* and the differentiation compartment as having a growth rate of *α_m_*. For the division symmetry fraction *ϕ*, we assume that it is kept equal across both compartments.

## V. ANALYTICAL SOLUTION OF THE TWO-COMPARTMENT MODEL

We then examined for which values of the growth rates of the two compartments, *α_n_* and *α_m_*, the system is in homeostasis and asked what the resulting steady state number of dividing cells *D* would be, given all parameters of the system.

We addressed this problem using an analytical approach. A trivial solution is *α_n_, α_m_* = 0, which would be equivalent to the uniform model with *α* = 0, as discussed above. Here, we will focus on all other possible solutions. For the derivation of the steady-state number of proliferating cells, we ignore that there is variation in the cell cycle time, and simply define the cell cycle time as *T*.

Using a transport model we arrive at the following two equations for the niche compartmant:

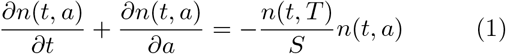

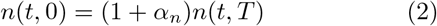

The full derivation can be found in Appendix A. In the above equations, *a* is the age of a cell, with *a* = 0 for newly-born cells and *a* = *T* at the moment of a cell division. *n*(*t, a*) is the number of cells at a given time with a given age, and integrating over all values of *a* results in *N*(*t*). The transport model describes the number of cells of age *a* to *a* + *da* at time *t*. This number changes due to aging of cells, cell divisions and due to proliferating cells exiting the niche compartment. At every cell division, on average 1+*α_n_* proliferating daughter is born. In addition, exactly one cell is ejected from the niche compartment, which has a chance of 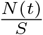 of being a proliferating cell.

For the differentation compartment, we arrive at similar equations (Eq. A5), except that the right hand side is now positive because cells enter the compartment instead of exiting. In steady state, we arrive at the following solution for the total number of dividing cells:

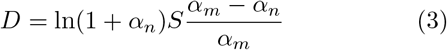

This equation is a central result of our study, and it has several implications.

1. The equation has solutions for *D* > 0 if *α_m_* < 0 and *α_n_* > 0. This dictates that the system can be in a steady state for other growth rates than *α_n_, α_m_* = 0. As long as the growth rate of the niche compartment is positive and the growth rate of the differentiation compartment is negative, a steady state is reached. For other values, the systems either decays or grows without bounds.
2. The number of proliferating cells in the entire system *D* increases as *α_n_* and *α_m_* increase. The number of proliferating cells *D* = 0 independent of *α_m_* for *α_n_* = 0, while it increases well beyond the niche size *S* for *α_m_* ≈ 0.
3. The equilibrium number of proliferating cells in the system is independent of both the cell cycle time *T* and the division symmetry fraction *ϕ*.
4. The number of proliferating cells in the entire system *D* always scales linearly with the total size of the niche compartment. There is no combination of growth rates *α_n_* and *α_m_* for which this linearity is lost.

## VI. STOCHASTIC SIMULATIONS OF THE TWO-COMPARTMENT MODEL

### A. Impact of fluctuations

From our analytical results, we have seen what values of the growth rates of the two compartments, *α_n_* and *α_m_*, result in a homeostatic system. However, it leaves open how sensitive this steady state is to fluctuations. For an uniform system, we saw that asymmetric divisions resulted in a system with lower fluctuations in the number of proliferating cells. We therefore asked if more division asymmetry also results in fluctuations in the two-compartment model.

We start by performing a parameter sweep, sampling all possible combinations of *α_n_, α_m_* and *ϕ*. For now, we keep the initial number of proliferating cells at *D*=30. The niche compartment size *S* is calculated using Eq. 3. As a result, *S* varies with both *α_n_* and *α_m_*. We reproduced the expected average value 〈*D*(*t*)〉=30 for all combinations of *ϕ*, *α_n_* and *α_m_* (Fig. S1), confirming the validity of our analytical result in Eq. 3.

In Fig. 3a, we show *D*(*t*) for two parameter sets that differ strongly in the degree of symmetry. In the first case, almost all cells divide asymmetrically, while *α_n_, α_m_* = 0, thus corresponding to the strategy that minimizes fluctuations in the uniform model. In the second, all cells divide symmetrically, while the growth rate differs strongly between compartments. Interestingly, we find that the parameter set with high fraction of symmetric divisions results in smaller fluctuations in *D*(*t*). This is in contrast with the uniform model, where asymmetric divisions always provided less fluctuations compared to symmetric divisions.

**FIG. 3.**
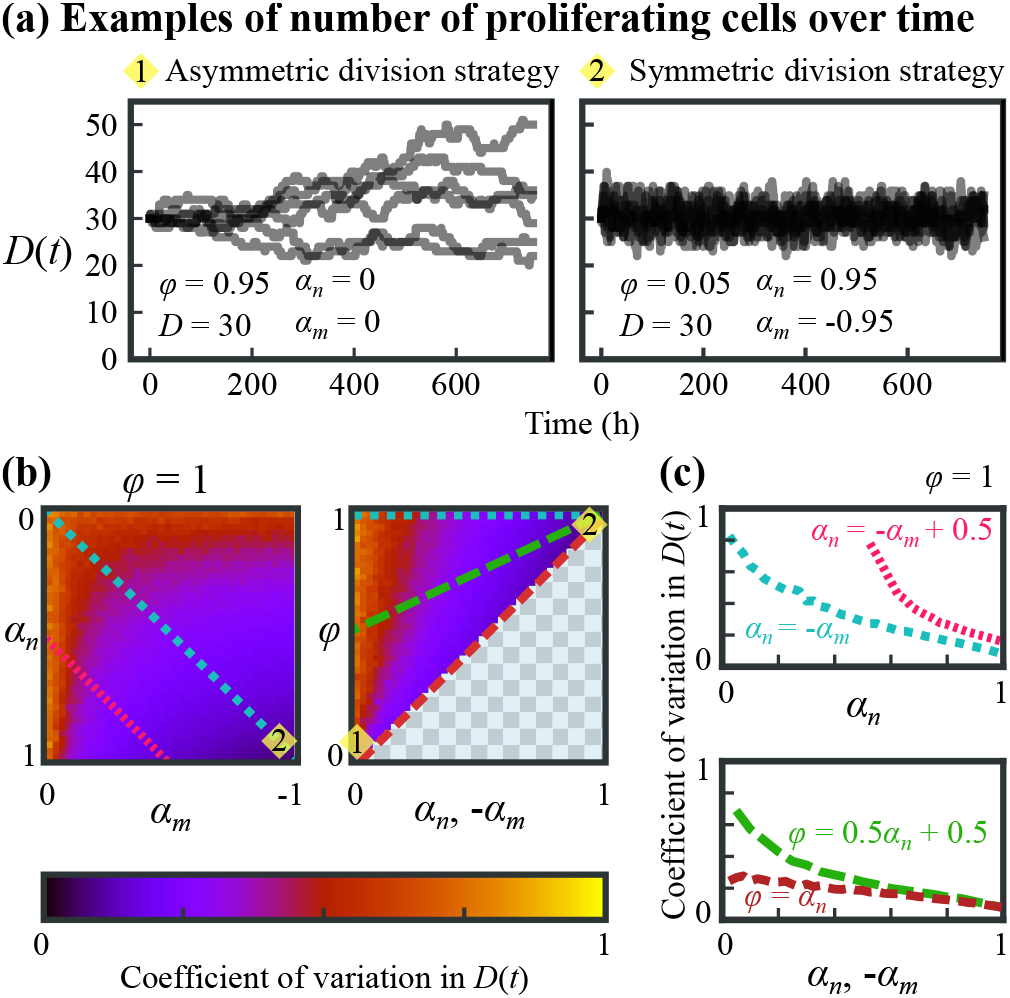
Exploration of the two-compartment model. Simulations ran 10^5^ hours for each parameter set. The initial number of dividing cells *D* = 30 and niche compartment size *S* is set according to Eq. 3. **(a)** Two panels showing *D*(*t*) for six example simulations each of the optimal parameters for either an asymmetric divisions strategy (left) or a symmetric division strategy (right). **(b)** Overview of the coefficient of variation in the number of dividing cells *D*(*t*). The blocked areas represent impossible combinations of parameters, such as strictly asymmetric divisions with a positive growth rate. **(c)** Coefficient of variation along selected lines. Line style matches the lines of panel b.

In (Fig. 3b) we quantify the coefficient of variation in *D*(*t*) for different combinations of *α_n_*, *α_m_* and *ϕ*. In general, we find that fluctuations decrease when the difference in growth rate between compartment, *α_n_ – α_m_*, increases. When we focus on the the subset of parameter sets with *α_n_* = – *α_m_*, we find that for a fixed value of the two growth rates, fluctuations are reduced by increasing the fraction of asymmetric divisions (decreasing *ϕ*), which appears consistent with our results for the uniform model and at odd with our observation in Fig. 3a that high symmetry resulted in low fluctuations.

To examine this further, we compare in Fig. 3c the coefficient of variation along the four lines shown in Fig. 3b. In the top panel, we compare two lines for parameter sets with only symmetric divisions (*ϕ* =1) that have either a larger (cyan) or smaller difference (pink) in growth rate between the two compartments. In both cases, the coefficient of variation in *D*(*t*) decreases with increasing difference in growth rates, but the parameter sets with the smaller difference (pink) always showed larger fluctuations.

The bottom panel shows two other lines, where we varying *ϕ* while fixing *α_n_* = – *α_m_*. The red line uses for every value of *α_n_* the lowest possible value of *ϕ*, i.e. the maximum amount of asymmetry that the growth rate permits. At *α_n_* = 1 all divisions are symmetric. The green line has a higher fraction of symmetric divisions. The lines show that for a given growth rate *α_n_*, the parameter set with the highest asymmetry results in the smallest coefficient of variation in *D*(*t*), as seen above. However, at the same time the global minimum of the coefficient of variation is found for *α_n_, –α_m_* = 1. This is because the decrease in fluctuations is strongest when difference in the two growth rates is maximal. For these values of the growth rate only *ϕ* = 1 is allowed, which is a system with only symmetric divisions. Therefore, although asymmetry generally reduces the coefficient of variation in *D*(*t*), the optimal solution is a fully symmetrically dividing system.

Based on Fig. 3b and c, we can conclude that the in the two compartment model the best strategy to minimize fluctuations in the number of proliferating cells *D*(*t*) is to use a system where all cells in the niche compartment proliferate, but each cell born outside the niche compartment immediately stops proliferating. This results in dominance of symmetric cell divisions. Other solutions, such as strictly asymmetric divisions or a combination of symmetric and asymmetric divisions result in more fluctuations in the number of proliferating cells.

### B. Dependence of fluctuations on niche size

Next, we wondered what the influence of the size of the niche compartment would be on the stability of the system. For each size, we plot the coefficient of variation along two lines. For the first line, we take the line that describes all points for which *ϕ* = *α_n_, – α_m_*, corresponding to the red dashed line in Fig. 3b. This is the line with the lowest coefficient of variability in the number of dividing cells *D*(*t*) for every *ϕ*. For the second line, we examine the dependence of the niche compartment size *S* for the parameters on the line with *ϕ* =1 and *α_n_* = – *α_n_*, corresponding to the cyan dotted line in Fig. 3b. This line represents the line with the largest difference in coefficient of variability of *D*(*t*); it goes from the global minimum to the global maximum of the coefficient of variability in *D*(*t*) for a given *S*.

The results are displayed in Fig. 4a. Consistent with the results above, for every given size, increasing the difference in growth rates *α_n_* and *α_m_* will always result in a system with a lower coefficient of variation in *D*(*t*) and a lower depletion rate. Moreover, at larger compartment sizes *S* the coefficient of variation in *D*(*t*) always becomes lower. Therefore, independent of the chosen parameter set, a system can always decrease its relative fluctuations by creating a larger niche. For the same reason, decreasing the niche compartment size results in more frequent depletions of the proliferating cells (Fig. 4b).

**FIG. 4.**
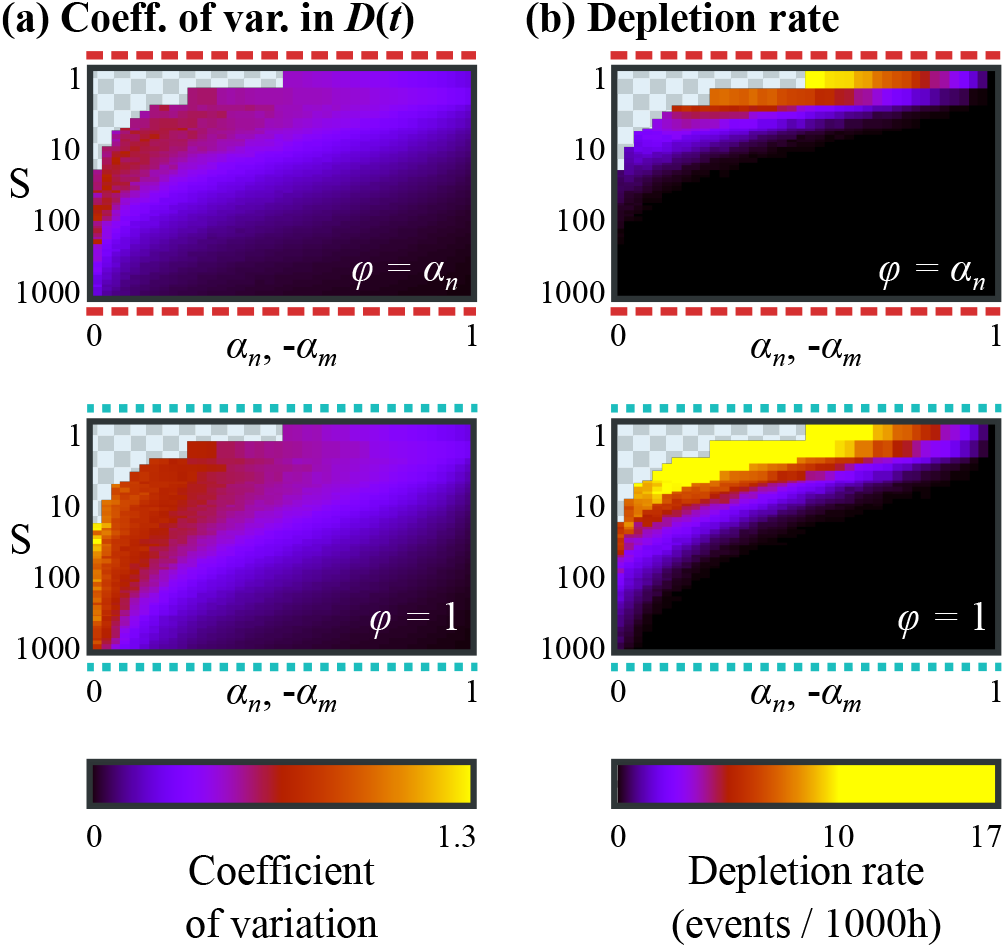
Simulations for different niche compartment sizes *S*, each for 10^5^ hours. The initial number of dividing cells *D* is set according to Eq. 3. The striped lines above and below the graphs correspond to the lines in Fig. 3. The blocked areas in the graphs are parameter ranges for which Eq. 3 does not give a solution. **(a)** Coefficient of variation of *D*(*t*) for various parameters. **(b)** Depletion rate for various parameters.

The observation that a smaller system results in more fluctuations is as expected. However, interestingly even for small niche sizes the system can remain stable, provided the symmetry fraction is high. Decreasing the niche compartment size from *S* = 30 to only *S* = 10 still results in a system with low fluctuations (less than 1 collapse per 10 years), provided the niche compartment maintains a high growth rate and symmetry fraction.

### C. Determining growth rate and division symmetry by clone size distributions

Experiments often measure clone size distributions using lineage tracing. For instance, using a model analogous to the uniform model above, the division symmetry *ϕ* was estimated by fitting to experimental long-term clone size scaling distributions, measured for different time frames [15]. We therefore asked whether parameters such as growth rate and division symmetry could also be inferred from clone size distributions obtained in the context of the two-compartment system.

For short-term clone size distributions, dominance of symmetric divisions appears clearly as an enrichment of even clone sizes. In Fig. 5a we show simulated clone size distributions of 1000 simulation runs, taken after 48 hours. For the case where symmetric divisions are dominant (*ϕ* = 0.95, displayed in Fig. 5a, left), we can see that even clone sizes 2, 4, 6 and 8 all occur in higher frequencies than the odd clone sizes 3, 5 and 7. This enrichment is not visible for systems where asymmetric divisions are dominant, or where neither division type is dominant (Fig. 5a, center and right).

**FIG. 5.**
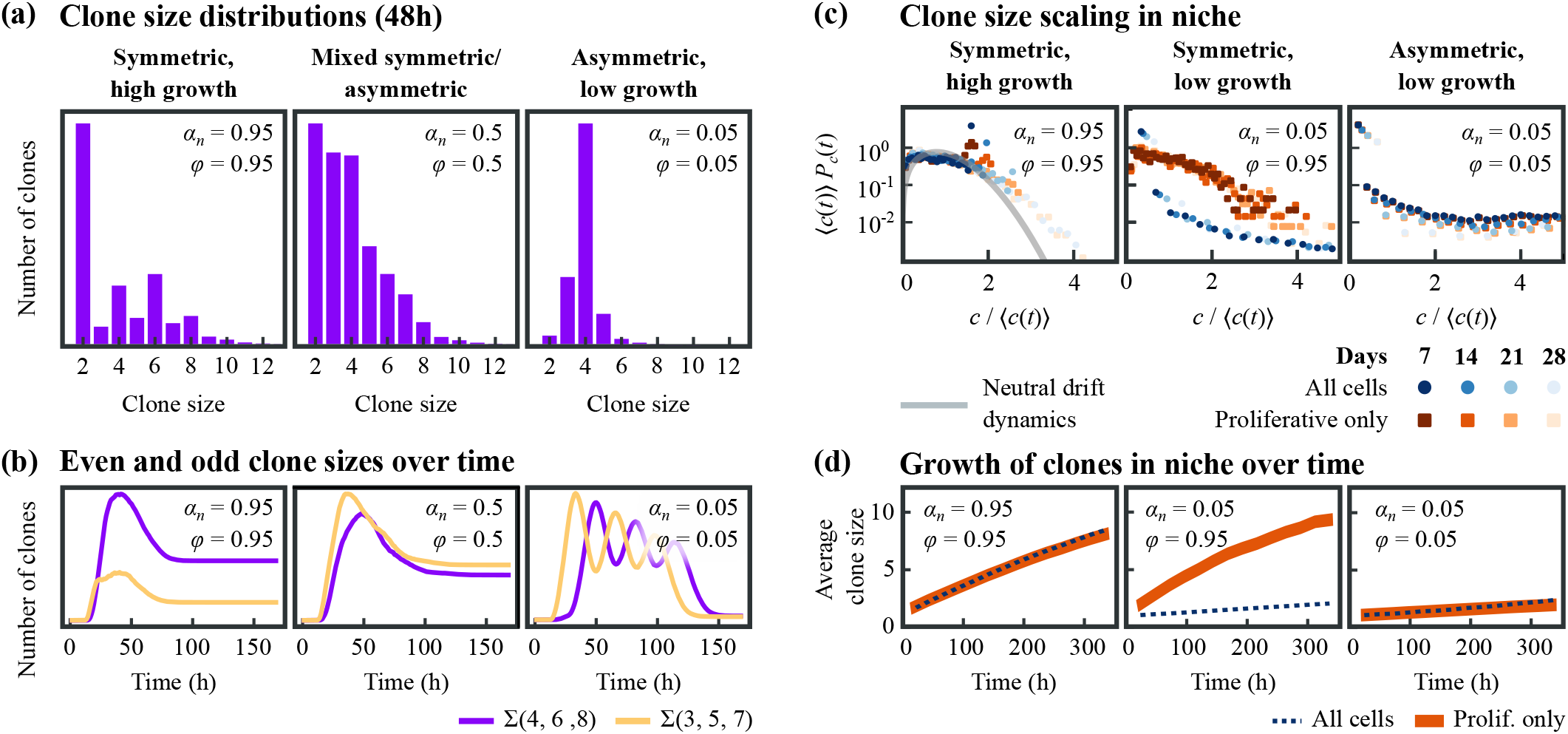
Clone size distributions of cells within the niche compartment. 1000 simulations, *α_m_* = – *α_n_, D* = 30 and *S* is set according to Eq. 3. **(a)** Short-term clone size distributions, taken after 40 hours. **(b)** Amounts of even and odd clone sizes over time. For the symmetric, high growth case there is initially a large enrichment of even clone sizes, which becomes smaller after 50 hours. For the mixed symmetric/asymmetric case, odd clone sizes are enriched. For the asymmetric case, all clones grow at almost the same rate, and therefore a single clone size is dominant at any point in time. This causes oscillatory behavior in whether even or odd clone sizes are dominant. **(c)** Clone size scaling for the symmetric, low growth case. For these parameters, there is a large contrast between the clone size distribution of all cells and the clone size distribution of only the proliferating cells. **(d)** Growth of the average clone size over time.

However, as the clone size over time in Fig. 5b shows, this enrichment of even clone sizes is strongest until 60 hours. In addition, during the entire simulation the enrichment is barely visible for clone sizes larger than 8 (Fig. 5a, Fig. S2). In the simulation this has two causes. First, we simulate for *ϕ* = 0.95, which still results in 5% of all divisions being asymmetric. While symmetric divisions will keep the clone size of a single lineage even, asymmetric divisions will not. Already a single asymmetric division anywhere in the lineage tree results in the clone size becoming odd.

Second, the variability in cell cycle times also contributes to the occurrences of odd clone sizes. Consider for example two proliferating sister cells. Initially the clone size is two, and after both sisters have divided it is four. However, in between the divisions of both sisters, the clone size is 3 and therefore odd. This effect occurs more often for large lineage trees, simply because there are more sister pairs, therefore increasing the chance of at least one of them making the clone size odd. In conclusion, the effect of division symmetry is visible only on the time scale of a few divisions.

Often, clone size growth is characterized by the clone size scaling function [5], which unlike the distributions discussed above are independent of time. Would there be a way to measure both the symmetry fraction and the growth rate from this scaling function? In other words: do the symmetry fraction and growth rate affect the observed long-time scaling behavior?

In experiments, clone size distributions are often collected only in the stem cell niche. Moreover, our simulation does not implement cell death, which results in the number of non-proliferating cells outside the niche compartment growing continuously. Such unbounded growth would prevent any long-term scaling behavior. For that reason, we will in our analysis ignore all cells outside the niche compartment.

We calculated the scaling function of different combinations of the symmetry fraction *ϕ* and growth rate *α_n_*. In Fig. 5c we show the scaling functions, both for all cells in the niche compartment and for only the proliferating cells. Interestingly, the shape of the scaling function for the proliferating cells in the niche is almost independent on the growth factor *α_n_*, as both for *α_n_* = 0.05 (Fig. 5c, left) and *α_n_* = 0.95 (Fig. 5c, center) the scaling function follows a concave shape. Instead, the scaling function depends on the symmetry fraction *ϕ*: for high symmetry fractions the scaling follows the concave pattern predicted by neutral drift dynamics [9] (Fig. 5c, left and center), while for low symmetry fractions the scaling function follows a convex shape (Fig. 5c, right).

However, if we look at all cells, then the scaling function no longer depends on the symmetry fraction *ϕ*, but on the niche growth rate *α_n_*. For high *α_n_* the scaling function follows a concave shape (Fig. 5c, left), while for low *α_n_* the function follows a convex shape (Fig. 5c, center and right). The scaling functions of other parameter sets are displayed in Fig. S3, and are consistent with these observations.

The cause of this scaling behavior can be seen in Fig. 5d. In this panel, we plot the average clone size over time for the same three parameter sets. As expected, the average clone size for all cells in the niche compartment increases faster for a high growth rate (Fig. 5d, left) compared to a low growth rate (Fig. 5d, center and right). However, if we include only the proliferating cells, then the clone size growth depends on *ϕ*. For high *ϕ* (Fig. 5d, left and center), the clones grow faster than for low *ϕ* (Fig. 5d, right). This is because the number of proliferating cells can only grow due to symmetric divisions, as asymmetric divisions do not have an effect on the number of proliferating cells. Even though the total number of proliferating cells is independent on *ϕ* (Eq. 3), the proliferating cells are now distributed over more clones. We can see that the fast-growing clones in Fig. 5d correspond to the concave scaling functions in Fig. 5c, and the slow-growing clones correspond to the convex scaling functions.

In conclusion, this scaling behavior in principle provides a way to experimentally determine the symmetry fraction *ϕ* and the growth rate *α_n_* using lineage tracing techniques that use a proliferation marker.

## VII. DISCUSSION

In this paper, we investigated the impact of fluctuations for different stem cell maintenance strategies, in the context of a stem cell niche. For this, we used a model with two compartments and two cell types, namely proliferating cells and non-proliferating cells. The model assumes a stem cell niche that is restricted in size, and that cells decide whether to continue proliferating or not depending only on the identity of the compartment in which they are born. No other interactions between cells are assumed, and besides their proliferation state, no other internal cellular state is considered.

The model predicts two possible strategies for minimizing stem cell number fluctuations under homeostatic conditions. If the two compartments do not differ in growth rate, then the growth rate must be zero to balance proliferation and cells must maximize the fraction of asymmetric divisions. This special case of our model corresponds to the model developed previously by Klein et al.[15], and also used by Sei et al.[21]. However, if the two compartments are allowed to differ in growth rate, then another strategy with lower cell number fluctuations becomes available. In this strategy, fluctuations are minimized when the different in growth rate between the two compartments is maximized and hence all divisions are symmetric. This finding is in contrast with the view that asymmetric divisions offer a more regulated stem cell maintenance process in stem cell niches [25, 26], but consistent with measurements performed recently by us that show that symmetric divisions dominate in growing intestinal organoids[24]. In this optimal limit, the dynamics in the niche compartment of our model becomes similar to the models of Snippert et al.[10] and Lopez-Garcia et al.[9]. Our results thus show that their models, which were based on experimental observations, correspond to a parameter regime, in our more general model, that minimizes fluctuations in the number of proliferating cells.

It would be interesting to experimentally test whether other organs or tissues that contain stem cell niches exhibit dynamics consistent with the optimal limit of our model, with strong difference in growth rate between the stem cell niche and the rest of the tissue, and most divisions symmetric, producing either two proliferating or two non-proliferating daughters. Our analysis shows that clone size distributions might be able to test this.

## ACKNOWLEDGMENTS

We would like to thank Max A. Betjes for useful discussions, suggestions and comments. R.K. was funded by an NWO Building Blocks of Life grant from the Dutch Research Council, number 737.016.009, https://www.nwo.nl/.

## Appendix A: Derivation of Eq. (3)

When a cell is born, it is set to be either proliferating or non-proliferating. Proliferating cells will divide *T* hours after they are born, while non-proliferating cells will not divide. Here, *T* is a normal distribution with We define *n*(*t, a*)*da* as the number of proliferating cells in the niche compartment at time *t* with age bracket (*a, a+da*). Here, age is defined as the time since the last division. The number of cells flowing in and out of this age bracket due to aging is:

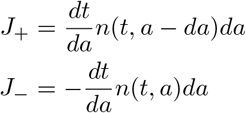

The rates for cells entering and exiting the niche compartment are defined as *k*_+_ (*a*) and *k*_−_(*a*), respectively. Note that in our simulation cells do not reenter the niche compartment so *k*_+_(*a*) = 0.

Using this information, we look at how the number of cells of a particular age bracket evolves over time. This number is equal to the number of existing cells of that age bracket, plus the number of incoming cells due to aging, minus the number of cells exiting the age bracket due to aging, plus the number of cell changes due to cells entering and exiting the compartment.

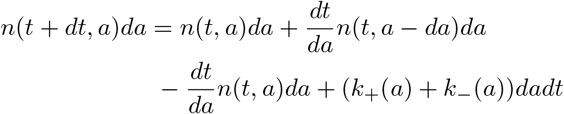

Rearranging, we find:

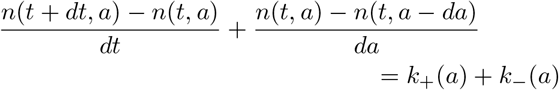

Using the definition of the derivative we readily find:

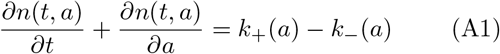

### 1. Growth boundary condition

Next, we introduce the growth boundary condition. If there are no cells moving in between compartments then *k*_+_(*a*) = *k*_−_ (*a*) = 0 and *dN* = *J_in_* – *J_out_*. Here, 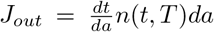 is the rate of cells exiting the cell cycle, and *J_in_* the rate of cells entering the cell cycle. As each division produces on average *α* +1 proliferating cells, *J_in_* = (*α* + 1) · *J_out_*. Together:

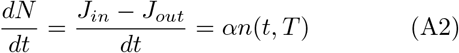

From the definition 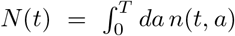 it follows that:

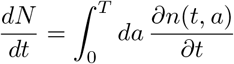

From Eq. A1 in the case where *k*_+_(*a*) = *k*_−_(*a*) = 0, we can see that 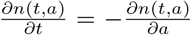. Inserting this in the above integral and evaluating it result in:

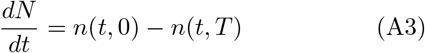

Combining Eq. A3 with Eq. A2 results in:

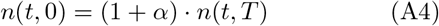

### 2. Full model

Having satisfied the growth condition, we will now calculate the values of *k*_−_(*a*) and *k*_+_(*a*), which represents the movement of cells between compartments. In our model, cells cannot re-enter the niche compartment, so *k*_+_(*a*) = 0. For expressing *k*_−_(*a*), we realize that the fixed size of the niche compartment means that for every cell division, one random cell must be removed from the niche compartment and added to the differentiation compartment. Therefore, *k*_−_(*a*) is equal to the number of cells dividing in a given time interval, multiplied by the chance that a cell being ejected is a dividing cell, multiplied by the probability that the ejected cell has the age (*a, a* + *da*):

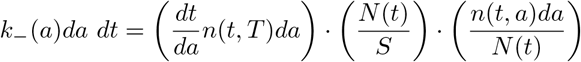

Here, we remind the reader that *N*(*t*) is defined as the total number of proliferating cells in the niche at time *t*, and that *n*(*t, a*) is the number of proliferating cells in the niche compartment of age *α* at time *t*.

From this equation it directly follows that:

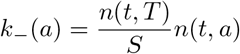

Inserting these results in Eq. A1, we obtain Eq. 1:

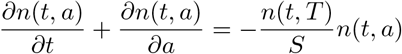

The number of newly-born proliferating cells in the niche was already defined by Eq. A4.

For the differentiation compartment a similar analysis can be made, noting that here *k*_−_ (*a*) is now negative, and *k*_+_(*a*) is now equal to *k*_−_(*a*) of the niche compartment. The number of proliferating cells in the differentiating compartment of age *a* at time *t* is defined as *m*(*t, a*).

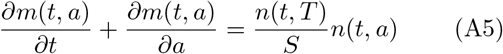

The equivalent of Eq. A4 for the differentiation compartment simply becomes:

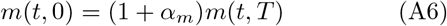

### 3. Steady state for the niche compartment

To solve the number of proliferating cells, we assume that the age distribution is exponential:

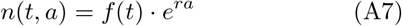

Here, *f*(*t*) is a function independent of *a* and *r* is a coefficient. At *a* = 0 we obtain *n*(*t, a* = 0) = *f*(*t*). From Eq. A4 we obtain *n*(*t, a* = 0) = (1 + *α_n_*) · *n*(*t,T*) = (1 + *α_n_*) · *f*(*t*)*e^rT^*, where we inserted Eq. A7. This results in:

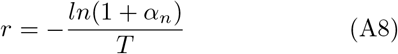

Next, we substitute Eq. A7 into Eq. 1:

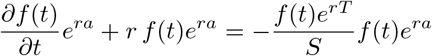

By dividing by *e^ra^* and rearranging, we find:

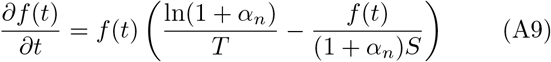

In steady state, *f*′ = *f*(*t*) so that 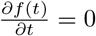. Therefore, from Eq. A9 two solutions arise. The first is the trivial solution *f*′ = 0, the second

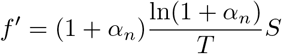

Inserting this solution into Eq. A7 (with Eq. A8) results in:

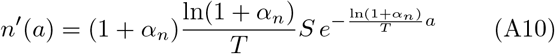

Integrating over all values of *a* results in the number of dividing cells in the niche compartment:

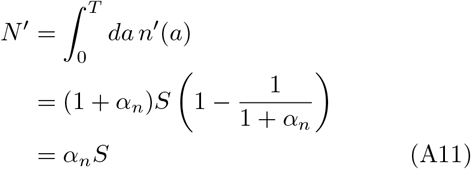

Here, we used:

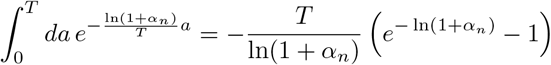

### 4. Steady state for the differentiation compartment

To obtain the total number of dividing cells, we also need to calculate the number of cells in the differentiation compartment. For this compartment, we need to take into account that there is an incoming supply of dividing cells from the niche compartment. In steady state, 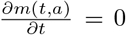. To ensure homeostasis in the full system, 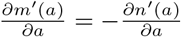. From Eq. A10 we therefore obtain:

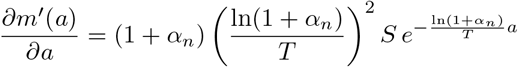

As the age distribution is exponential, we can state:

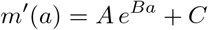

From inserting this equation into the one above it and comparing terms, it follows that 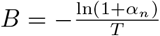 and 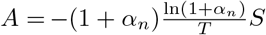.

From Eq. A6 we obtain *m*′(0) = (1 + *α_m_*)*m*′(*T*), resulting in:

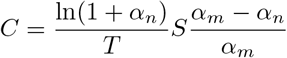

Together,

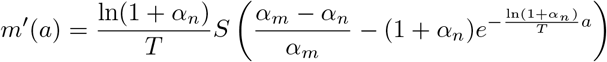

With 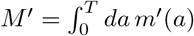 we obtain

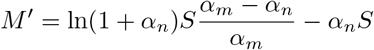

With *D* = *M*′ + *N*′ and the insertion of Eq. A11 we obtain the final result:

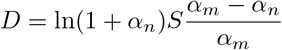

This equation has solutions for *D* > 0 if *α_m_* < 0 and *α_n_* > 0. If *α_n_* < 0 then *N*′ would be negative, and if *α_m_* > 0 then *M*′ would be negative, which are both not allowed.

## Appendix B: Average number of proliferating cells

**FIG. S1.**
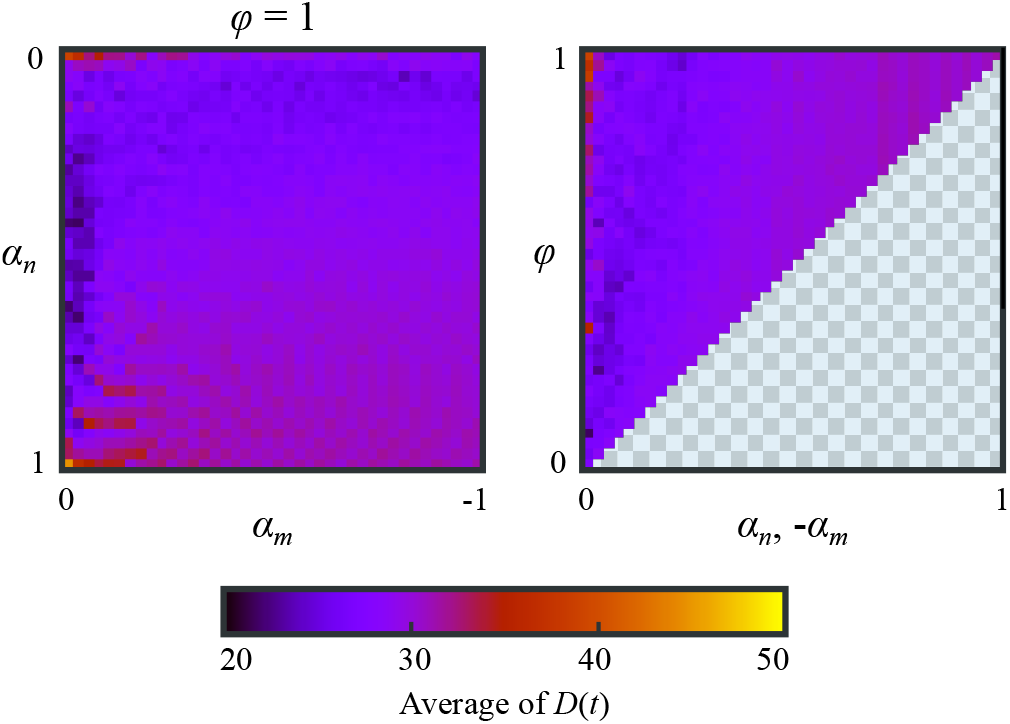
Average of *D*(*t*) for a 10^5^h simulations of each parameter set. The initial *D* was set according to Eq. 3. The figure uses the same layout as panel **(b)** of Fig. 3.

## Appendix C: Frequency of even and odd clone sizes over time

**FIG. S2.**
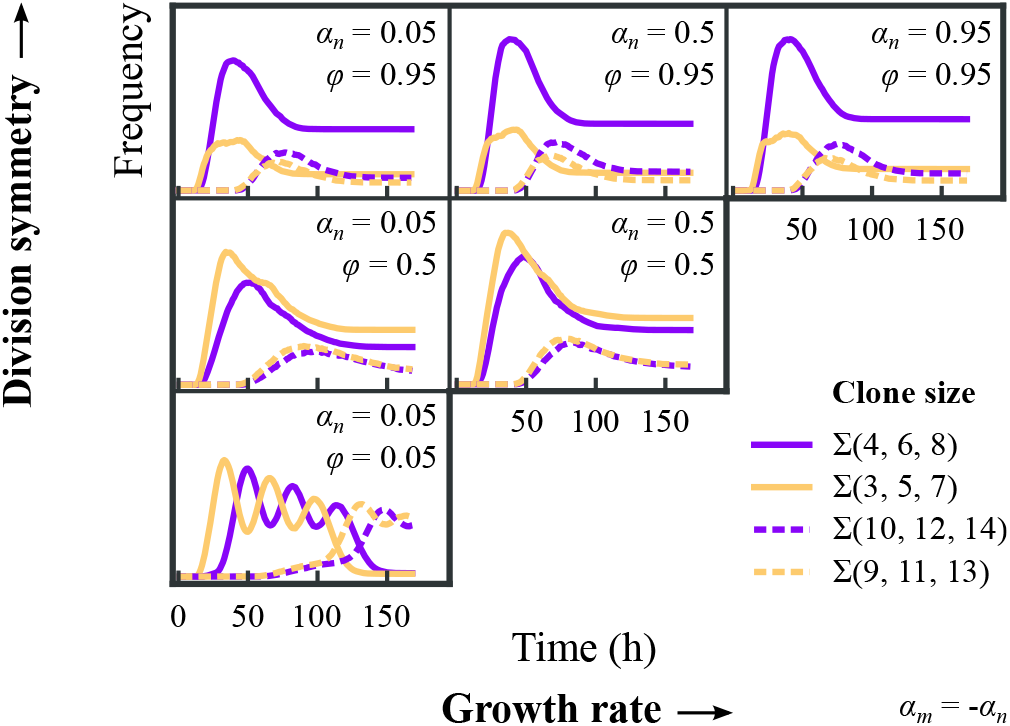
Clone sizes plotted over time for the entire system, for the sums of the given even and odd clone sizes. The evolution of the clone size depends mainly on the division symmetry, not on the growth rate.

## Appendix D: Clone size scaling functions

**FIG. S3.**
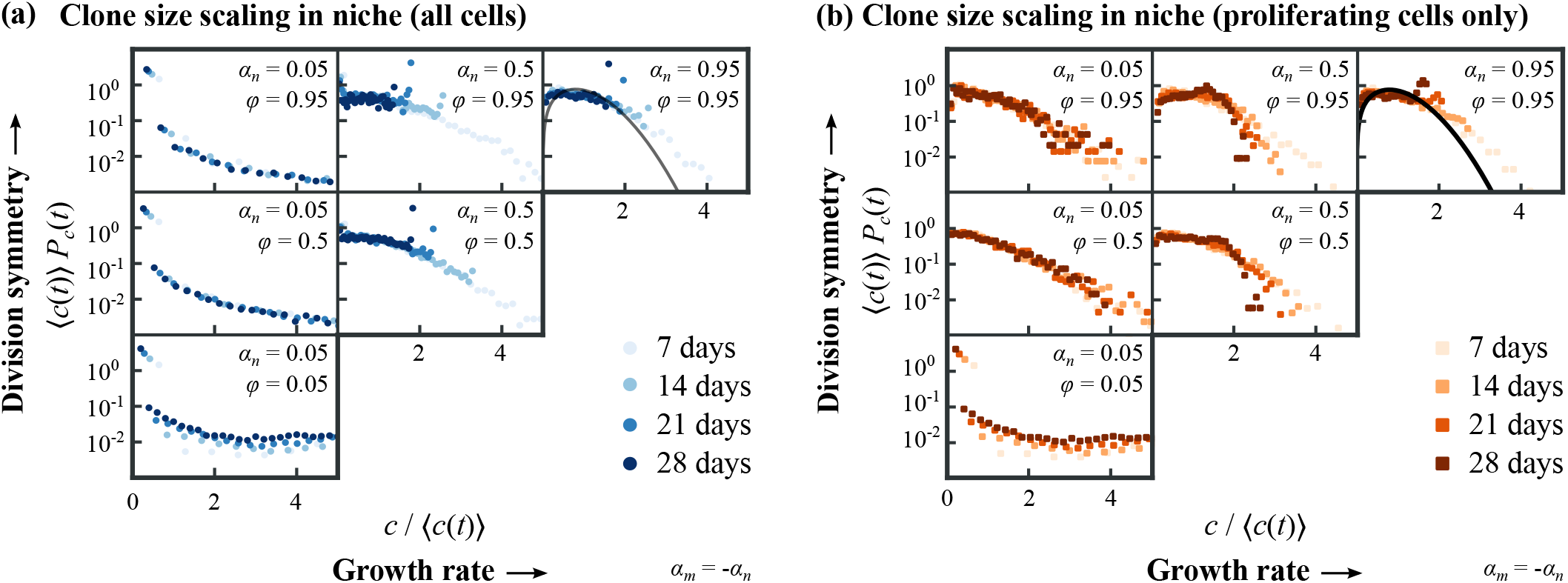
Scaling function for various configurations. **(a)** Scaling functions that include all cells in the niche compartment. **(b)** Scaling functions that include only the proliferating cells in the niche compartment.

## Notes

### Competing Interest Statement

The authors have declared no competing interest.

## References

[1] N. McCarthy, J. Kraiczy, and R. A. Shivdasani, Cellular and molecular architecture of the intestinal stem cell niche, Nature Cell Biology 22, 1033 (2020).

[2] S. D. C. Weterings, M. J. van Oostrom, and K. F. Sonnen, Building bridges between fields: bringing together development and homeostasis, Development 148, dev193268 (2021).

[3] B. D. Simons and H. Clevers, Strategies for homeostatic stem cell self-renewal in adult tissues, Cell 145, 851 (2011).

[4] A. Jilkine, Mathematical Models of Stem Cell Differentiation and Dedifferentiation, Current Stem Cell Reports 5, 66 (2019).

[5] L. Chatzeli and B. D. Simons, Tracing the Dynamics of Stem Cell Fate, Cold Spring Harbor Perspectives in Biology 12, a036202 (2020).

[6] A. M. Klein and B. D. Simons, Universal patterns of stem cell fate in cycling adult tissues, Development 138, 3103 (2011).

[7] Y. Kitadate, D. J. Jörg, M. Tokue, A. Maruyama, R. Ichikawa, S. Tsuchiya, E. Segi-Nishida, T. Nakagawa, A. Uchida, C. Kimura-Yoshida, S. Mizuno, F. Sugiyama, T. Azami, M. Ema, C. Noda, S. Kobayashi, I. Matsuo, Y. Kanai, T. Nagasawa, Y. Sugimoto, S. Takahashi, B. D. Simons, and S. Yoshida, Competition for Mitogens Regulates Spermatogenic Stem Cell Homeostasis in an Open Niche, Cell Stem Cell 24, 79 (2019).

[8] E. Clayton, D. P. Doupé, A. M. Klein, D. J. Winton, B. D. Simons, and P. H. Jones, A single type of progenitor cell maintains normal epidermis, Nature 446, 185 (2007).

[9] C. Lopez-Garcia, A. M. Klein, B. D. Simons, and D. J. Winton, Intestinal Stem Cell Replacement Follows a Pattern of Neutral Drift, Science 330, 822 (2010).

[10] H. J. Snippert, L. G. van der Flier, T. Sato, J. H. van Es, M. van den Born, C. Kroon-Veenboer, N. Barker, A. M. Klein, J. van Rheenen, B. D. Simons, and H. Clevers, Intestinal crypt homeostasis results from neutral competition between symmetrically dividing Lgr5 stem cells, Cell 143, 134 (2010).

[11] L. Ritsma, S. I. Ellenbroek, A. Zomer, H. J. Snippert, F. J. De Sauvage, B. D. Simons, H. C. Clevers, and J. Van Rheenen, Intestinal crypt homeostasis revealed at singlestem-cell level by in vivo live imaging, Nature 507, 362 (2014).

[12] H. Gehart and H. Clevers, Tales from the crypt: new insights into intestinal stem cells, Nature Reviews Gastroenterology & Hepatology 16, 19 (2019).

[13] H. Clevers and F. M. Watt, Defining Adult Stem Cells by Function, not by Phenotype, Annual Review of Biochemistry 87, 1015 (2018).

[14] P. Greulich, B. D. MacArthur, C. Parigini, and R. J. Sánchez-García, Universal principles of lineage architecture and stem cell identity in renewing tissues, Development 148, dev194399 (2021).

[15] A. M. Klein, D. P. Doupé, P. H. Jones, and B. D. Simons, Kinetics of cell division in epidermal maintenance, Physical Review E 76, 021910 (2007).

[16] Z. Sun and N. L. Komarova, Stochastic modeling of stemcell dynamics with control, Mathematical Biosciences 240, 231 (2012).

[17] B. Corominas-Murtra, C. L. Scheele, K. Kishi, S. I. Ellenbroek, B. D. Simons, J. Van Rheenen, and E. Hannezo, Stem cell lineage survival as a noisy competition for niche access, Proceedings of the National Academy of Sciences of the United States of America 117, 16969 (2020).

[18] V. L. Cannataro, S. A. Mckinley, and C. M. St. Mary, The implications of small stem cell niche sizes and the distribution of fitness effects of new mutations in aging and tumorigenesis, Evolutionary Applications 9, 565 (2016).

[19] V. L. Cannataro, S. A. McKinley, and C. M. St. Mary, The evolutionary trade-off between stem cell niche size, aging, and tumorigenesis, Evolutionary Applications 10, 590 (2017).

[20] L. Shahriyari and N. L. Komarova, The role of the bicompartmental stem cell niche in delaying cancer, Physical Biology 12, 055001 (2015).

[21] Y. Sei, J. Feng, C. C. Chow, and S. A. Wank, Asymmetric cell division-dominant neutral drift model for normal intestinal stem cell homeostasis, American Journal of Physiology Gastrointestinal Liver Physiology 316, G64 (2019).

[22] J. Yang, M. V. Plikus, and N. L. Komarova, The Role of Symmetric Stem Cell Divisions in Tissue Homeostasis, PLoS Computational Biology 11, 1 (2015).

[23] A. Azzalini and A. Capitanio, Statistical applications of the multivariate skew normal distribution, Journal of the Royal Statistical Society. Series B: Statistical Methodology 61, 579 (1999), arXiv:0911.2093.

[24] G. Huelsz-Prince, R. N. U. Kok, Y. J. Goos, L. Bruens, X. Zheng, S. I. Ellenbroek, J. van Rheenen, S. J. Tans, and J. S. van Zon, In preparation (2022).

[25] M. Inaba and Y. M. Yamashita, Asymmetric Stem Cell Division: Precision for Robustness, Cell Stem Cell 11, 461 (2012).

[26] Z. G. Venkei and Y. M. Yamashita, Emerging mechanisms of asymmetric stem cell division, Journal of Cell Biology 217, 3785 (2018).

